# Combinatorial interactions of *Hox* genes establish appendage diversity of the amphipod crustacean *Parhyale hawaiensis*

**DOI:** 10.1101/2022.03.25.485717

**Authors:** Erin Jarvis Alberstat, Kevin Chung, Dennis A Sun, Shagnik Ray, Nipam H. Patel

## Abstract

*Hox* genes establish regional identity along the anterior-posterior axis in diverse animals. Changes in *Hox* expression can induce striking homeotic transformations, where one region of the body is transformed into another. Previous work in *Drosophila* has demonstrated that *Hox* cross-regulatory interactions are crucial for maintaining proper *Hox* expression. One major mechanism is the phenomenon of “posterior prevalence”, wherein anterior *Hox* genes are repressed by more posterior *Hox* genes. Loss of posterior *Hox* expression under this model would predict posterior-to-anterior transformations, as is frequently observed in *Drosophila*. While posterior prevalence is thought to occur in many animals, studies of such *Hox* cross-regulation have focused on a limited number of organisms. In this paper, we examine the cross-regulatory interactions of three *Hox* genes, *Ultrabithorax (Ubx), abdominal-A (abd-A)*, and *Abdominal-B (Abd-B)* in patterning thoracic and abdominal appendages in the amphipod crustacean *Parhyale hawaiensis*. Studies of *Hox* function in *Parhyale* have previously revealed two striking phenotypes which differed markedly from what a “posterior prevalence” model would predict, including non-contiguous and anterior-to-posterior transformations. We probe the logic of *Parhyale Hox* cross-regulation by using CRISPR/Cas9 to systematically examine all combinations of *Ubx, abd-A*, and *Abd-B* loss of function in *Parhyale*. By analyzing homeotic phenotypes and examining the expression of additional *Hox* genes, we reveal *Hox* cross-regulatory interactions in *Parhyale*. From these data, we also demonstrate that some *Parhyale Hox* genes function combinatorially to specify posterior limb identity, rather than abiding by a posterior prevalence mechanism. These results provide evidence that combinatorial *Hox* interactions may be responsible for the tremendous body plan diversity of crustaceans.

**Graphical Abstract:** 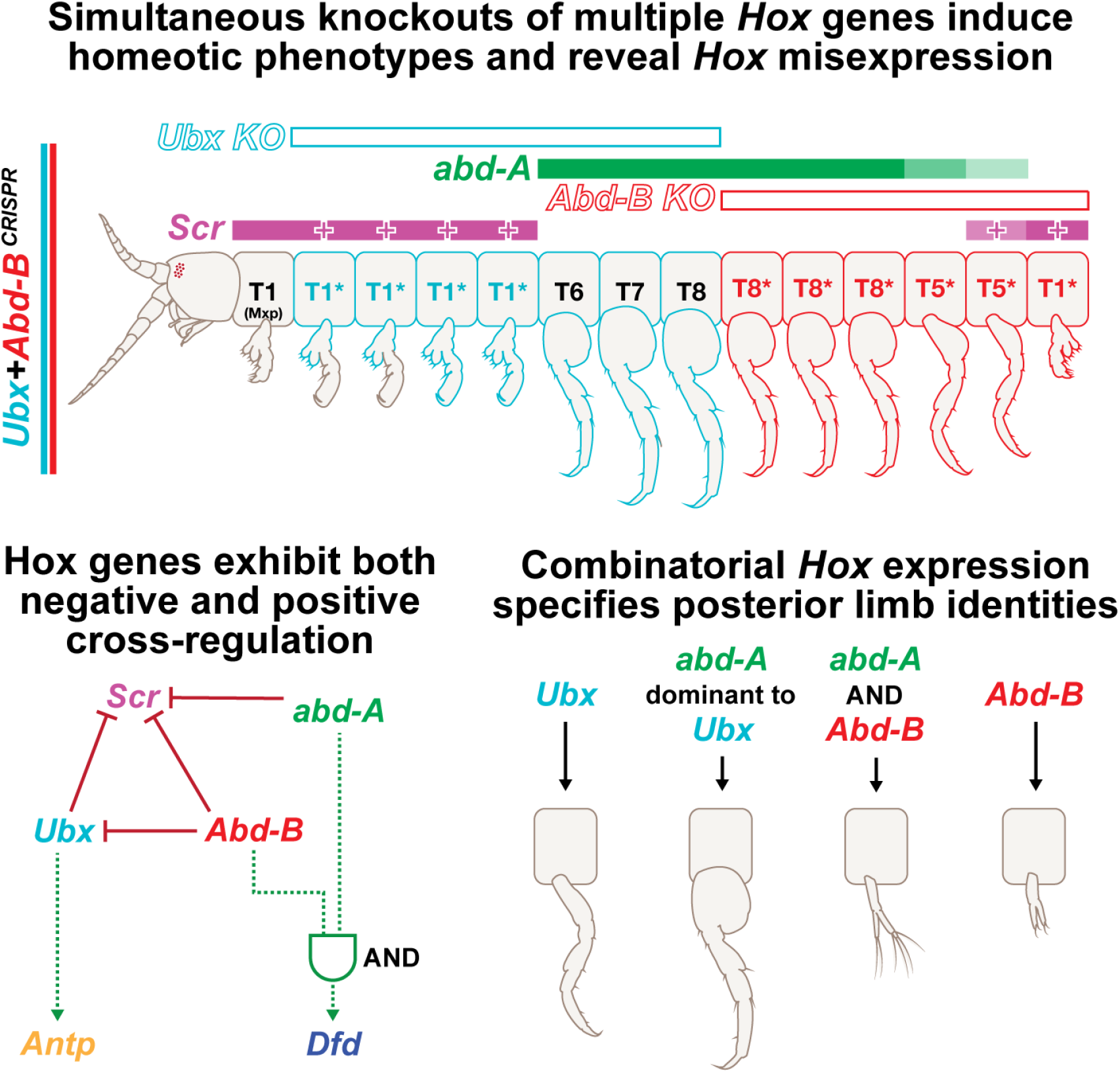

## Introduction

The *Hox* family of homeodomain-encoding transcription factors specifies regional identity along the anterior-posterior axis in most metazoans. These genes were first identified by their striking homeotic phenotypes, in which alterations of *Hox* expression or function caused transformations of one body part into another. For example, in the fruit fly *Drosophila melanogaster*, an *Antennapedia* gain-of-function (GOF) mutation can cause homeotic transformation of antennae into legs, while *Ultrabithorax* loss-of-function (LOF) causes homeotic transformation of halteres towards wing identity (Gehring et al., 1994; Graba et al., 1997; Mann and Morata, 2000; McGinnis and Krumlauf, 1992).

*Hox* genes are usually located in genomic clusters and have a conserved organization of paralogs that is mirrored by their expression along the anterior-posterior (AP) axis (Akam, 1989; Hughes and Kaufman, 2002; Krumlauf, 1994). *Hox* gene expression domains have distinct anterior and posterior boundaries and, in general, the posterior expression of one *Hox* gene partially overlaps the anterior expression of the neighboring and more posteriorly expressed *Hox* gene in the cluster. This ordered regional expression of *Hox* genes is widely conserved and acts to specify different morphologies through the activation and repression of distinct sets of downstream targets (Graba et al., 1997). This regionalization is reinforced by the refinement of *Hox* expression boundaries by neighboring *Hox* genes through cross-regulatory interactions, and through functional competition in the regions where the expression domains of multiple *Hox* genes overlap (Hughes and Kaufman, 2002; Levine and Harding, 1989).

The extensive conservation of *Hox* expression and regulatory interactions have led to a number of useful generalizations regarding the roles of *Hox* genes in body patterning (e.g., *Tribolium* and *Drosophila* (Reviewed in (Hughes and Kaufman, 2002; Shippy et al., 2008)). One key model called “posterior prevalence” has emerged from a clear AP directionality in the logic of homeotic transformations across different model organisms. In several arthropods, for example, studies have found that LOF of a posteriorly expressed *Hox* gene frequently results in homeotic transformation to the next-most anterior identity (a posterior-to-anterior transformation), while GOF mutations result in the opposite: anterior-to-posterior transformations (Reviewed by (Akam, 1998)). This phenomenon is related to the concept of phenotypic suppression, wherein more posteriorly expressed *Hox* genes are phenotypically dominant to more anteriorly expressed *Hox* genes (Macías and Morata, 1996; Noro et al., 2011).

Given their importance in establishing regional identity, *Hox* genes have been studied across taxa as potential drivers of body plan evolution. Increasingly, new technologies have enabled these investigations to include a greater diversity of species at greater functional detail. One recent example is the amphipod crustacean *Parhyale hawaiensis*, which boasts an abundance of diverse appendage types, relevant phylogenetic positioning, and a rapidly expanding collection of tools and genetic resources (Browne et al., 2005; Kao et al., 2016; Rehm et al., 2009; Sun et al., 2021). A complete analysis of *Hox* expression in *Parhyale* has revealed that each appendage type expresses a unique set of *Hox* genes (Serano et al., 2016), suggesting that a combinatorial “*Hox* code” may be responsible for establishing limb identity. Using CRISPR/Cas9 mutagenesis and RNAi, researchers have also examined the effects of ablating each *Hox* gene individually, revealing that the precise role of each *Parhyale Hox* gene in establishing regional identity follows the predictions made by examining expression domains (Liubicich et al., 2009; Martin et al., 2015). For example, knockout or knockdown of the *Hox* gene *Ubx* – normally expressed in the claws (chelipeds, T2/T3), forward walking legs (forward pereopods, T4/T5), and reverse jumping legs (reverse pereopods, T6-T8) – results in a loss of the T4/T5 identity and homeotic transformations of those appendages to more anterior limb types (Fig. 1B). These results reveal that *Ubx* is necessary for establishing forward walking leg identity.

**Fig. 1:**
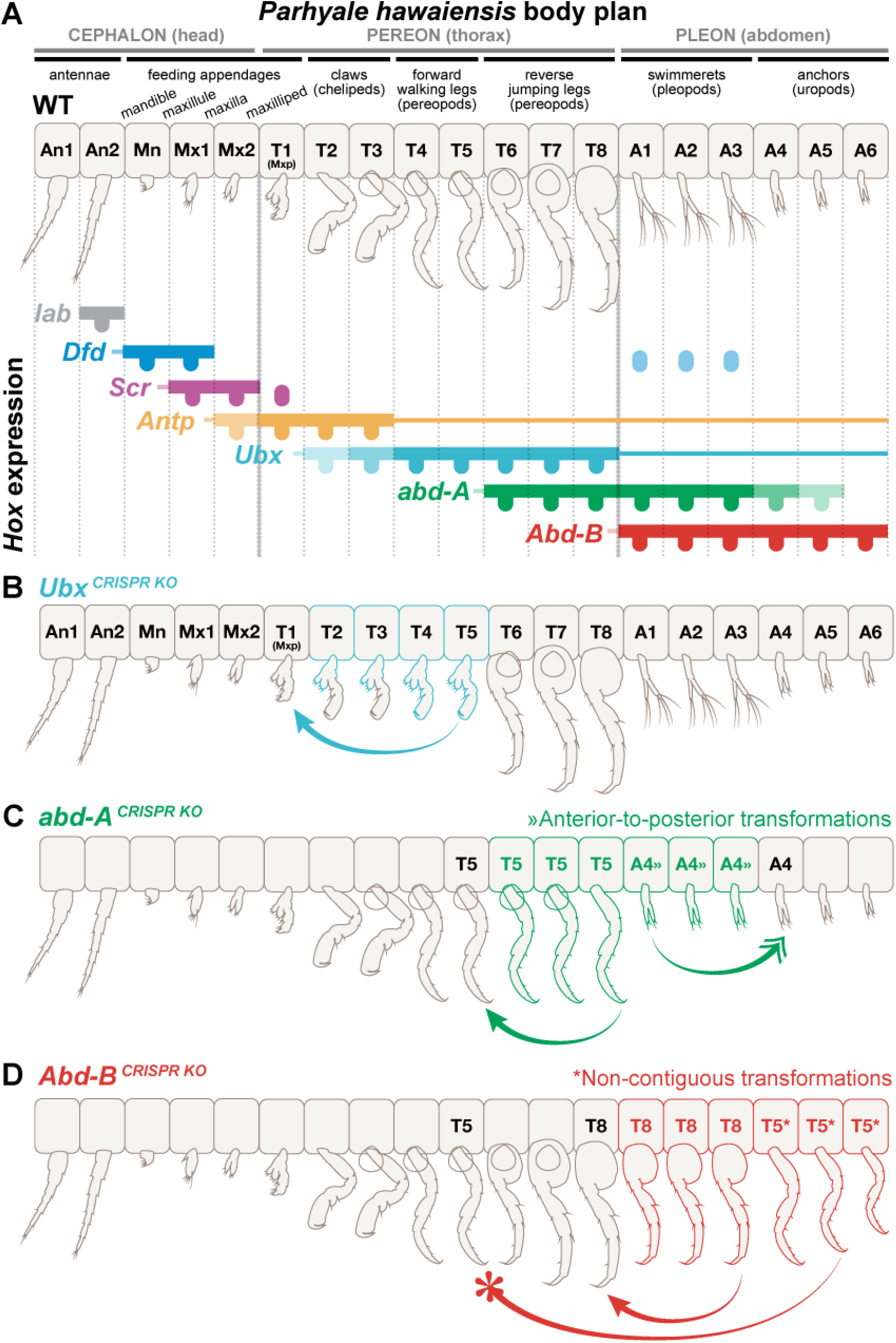
Schematic representation of Hox expression domains and corresponding limb morphologies in Parhyale. A) *Hox* gene expression in wildtype Parhyale appendages (Adapted from Serano et al. 2016). Expression in the body is marked by solid horizontal lines, and expression in the appendages is marked by ovals. Expression in the nervous system is marked by thin horizontal lines. Intensity of color reflects relative levels of expression; *Dfd* expression is also found in portions of the swimming legs (A1-A3). Expression not shown for *Hox* genes *pb* and *Hox3*, which have very limited domains spanning An2 and Mn, and do not include the entire appendage. Note that each unique combination of Hox genes corresponds roughly to a distinct appendage type: An1 (no Hox expression), An2 (lab, not shown), Mn (Dfd), Mx1 (Dfd + Scr), Mx2 (Scr + low levels of Antp), T1 Mxp (Scr + Antp), T2 - T3 Claws (Ubx + Antp), T4 - T5 walking legs (Ubx), T6 - T8 jumping legs (Ubx + abd-A), A1 - A3 pleopods (abd-A + Abd-B), and A4 - A6 uropods (Abd-B). Thoracic legs and abdominal appendages are patterned by the posterior Hox genes Ubx, abd-A, and Abd-B, corresponding to the Drosophila Bithorax complex. The expression domains of Ubx (expressed throughout the thorax excluding the first head-fused segment/maxillary appendage T1) and Abd-B (expressed throughout the abdominal) remain distinct from one another, while the expression domain of abd-A partially overlaps both, spanning the posterior thorax (where its anterior boundary delineates the T4 - T5 walking legs from the T6 - T8 jumping legs in the Ubx-expressing thorax) through anterior abdomen (where its posterior boundary delineates the A1 - A3 pleopods from the A4 - A6 uropods in the Abd-B-expressing abdomen. (B - C) Homeotic transformation by CRISPR/Cas9 Hox gene knockout (adapted from Martin et al. 2016). Arrows show direction of transformation, and color-outlined segments indicate which appendages are transformed. B) Abd-B knockout results in the transformation of the A1 - A3 pleopods to the T6 - T8 jumping leg phenotype that is immediately anterior as well as the non-linear skip in transformed identity of the A4 - A6 uropods towards that of the non-adjacent walking legs of T4 - T5. C) In the Ubx expressing thorax, abd-A knockout results in the transformation of the abd-A-expressing T6 - T8 jumping legs to the adjacently anterior T4 - T6 walking legs that do not express abd-A. The abd-A-expressing A1 - A3 swimming appendages are transformed to the adjacently posterior A4 - A6 uropods that fall outside the strong abd-A expression domain within the Abd-B-expressing abdomen. D) Ubx knockout results in the linear transformation of the thoracic legs T2 - T5 toward the maxillary identity of T1 that is adjacently anterior. Note that while all appendages expressing either abd-A or Abd-B are affected by the respective Hox gene knockout, only Ubx appendages not also co-expressing the more posterior abd-A are transformed upon Ubx knockout, as is consistent with the more traditional Hox models.

In addition to revealing the function of each individual *Hox* gene in establishing regional identity, this study also revealed a number of appendage transformations that were not easily explained by previous models of *Hox* function and regulation. Notably, knockouts of *Abd-B* and *abd-A* induced homeotic transformations that appeared to violate “posterior prevalence.” Knockout of *Abd-B*, which is expressed throughout the Parhyale abdomen, resulted in a non-contiguous “skip” in the transformation of the anchoring legs (uropods, A4-A6) toward forward walking legs (T4/T5)(Figure 1C). Additionally, knockout of *abd-A* induced both posterior-to-anterior transformations of the T6-T8 reverse jumping leg appendages to T4/T5 forward walking leg identity, as well as posterior-to-anterior transformations of swimming legs (pereopods, A1-A3) to A4-A6 anchoring leg identity (Figure 1D). Although these patterns of transformation are not predicted by classic invertebrate *Hox* logic, they can be logically explained using a combinatorial “*Hox* code” model, where the combined expression of multiple *Hox* genes would have combinatorial influence on downstream targets, rather than the more posterior *Hox* gene phenotypically dominating the establishment of limb identity (Kmita and Duboule, 2003).

Similar patterns of homeotic transformations more consistent with a “*Hox* code” model than with posterior prevalence have been described in other organisms. For example, mutation of the *Drosophila Hox* gene *Sex combs reduced (Scr)* in results in a bi-directional transformation similar in polarity to that observed in Parhyale *abd-A* KO (Duncan and Kaufman, 1975). However, the diverse array of different limbs found across the *Parhyale* body plan allow for a more comprehensive evaluation of this model.

In this study, we investigated these non-canonical transformations and the potential combinatorial nature of *Parhyale Hox* function using CRISPR/Cas9 mutagenesis. We generated knockout embryos for each paired combination of the posterior Hox genes *Ubx, abd-A*, and *Abd-B* (*abd-A* + *Abd-B, Ubx* + *abd-A*, and *Ubx* + *Abd-B*), as well as the triple knockout of all three genes (*Ubx* + *abd-A* + *Abd-B*). In addition to characterizing the resulting homeotic transformation phenotypes, we also performed comprehensive post-knockout expression analyses of other *Hox* genes to determine which *Hox* genes specify the newly-transformed identities we observed. These results reveal that *Hox* genes in *Parhyale* utilize both combinatorial function and cross-regulation to specify limb identity across the body plan.

## Results

### Exploring the ‘*Hox* code’ of thoracic and abdominal limb identities in Parhyale

#### *Abd-B* KO derepresses *Ubx* expression into the abdomen and induces abdomen-to-thoracic transformations

In *Abd-B* knockout hatchlings, A4-A6 anchoring legs are non-contiguously transformed toward T4/T5 forward walking leg identities, rather than to the T6-T8 reverse jumping leg identity, as would be predicted based on the “posterior prevalence” model (Fig. 1D). Previous work has demonstrated that the T4/T5 forward walking leg identity is regulated by the expression of the *Hox* gene *Ubx*. Loss of *Ubx* function using RNAi or CRISPR induces a loss of T4/T5 identities, whereas heat-shock induced overexpression of *Ubx* can cause transformations of anterior segments towards a T4/T5 walking leg identity (Liubicich et al., 2009; Martin et al., 2015; Pavlopoulos et al., 2009). We hypothesized that the non-contiguous transformation of A4-A6 to T4/T5 identity could be induced by expansion of *Ubx* expression to the posterior of the embryo as a result of loss of *Abd-B* expression.

To assess *Ubx* expression in *Abd-B* knockout embryos, we used a rat anti-*Parhyale Ubx* antibody (Liubicich et al., 2009). We observed that loss of *Abd-B* does result in a posterior expansion of *Ubx* into the abdomen of the animal, thus explaining the non-contiguous transformation previously observed (Fig. 2). This result suggests that *Abd-B* represses *Ubx* in *Parhyale*, a result that is consistent with a “posterior prevalence” relationship between these two *Hox* genes. However, additional examination of knockout embryos reveals that developing a full suite of posterior limbs of *Parhyale* requires combinatorial *Hox* function.

**Fig. 2:**
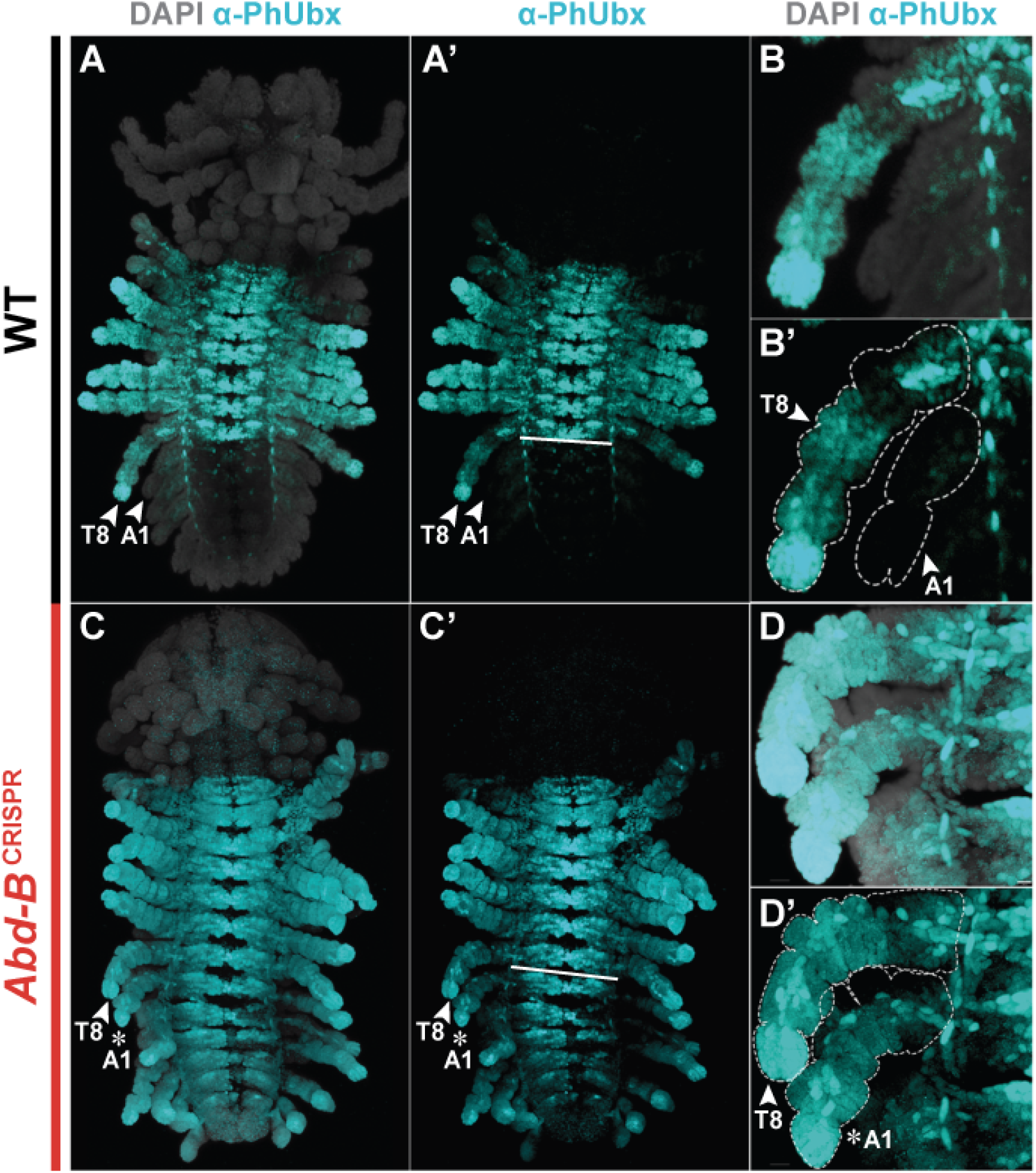
*Abd-B* KO induces posterior expansion of the Ubx boundary. (A-D’) Stage 22 Parhyale embryos stained for Ph-Ubx (Cyan) and counterstained with DAPI (gray). Arrowhead marks wildtype appendages; Asterisk denotes transformed appendages. Solid line marks the boundary between abdomen and thorax in A’ and C’. (A-B’) Ubx is expressed at high levels throughout the T4-T8 thoracic legs and at lower levels in the T2 - T3 claws. While Ubx appears to be expressed in select abdominal neurons/neuromeres, Ubx is absent from abdominal appendages. The thoracic leg vs. abdominal appendage phenotypes are clearly distinguishable in embryos at this stage, as is highlighted by the nascent T8 leg and A1 pleopod in (B-B’). Walking and jumping legs are uniramous with 7 podomeres (leg segments) while pleopods and uropods are biramous, with two rami (an endopod and exopod) emerging from a single protopod. (C-D’) *Abd-B* KO embryos show clear transformation of their abdominal appendages to a thoracic leg identity (D’ highlights uniramous, multi-podomeres of transformed A1). The posterior boundary of Ubx is very clearly expanded throughout the abdominal segments and appendages in the absence of *Abd-B*.

**Fig. 3:**
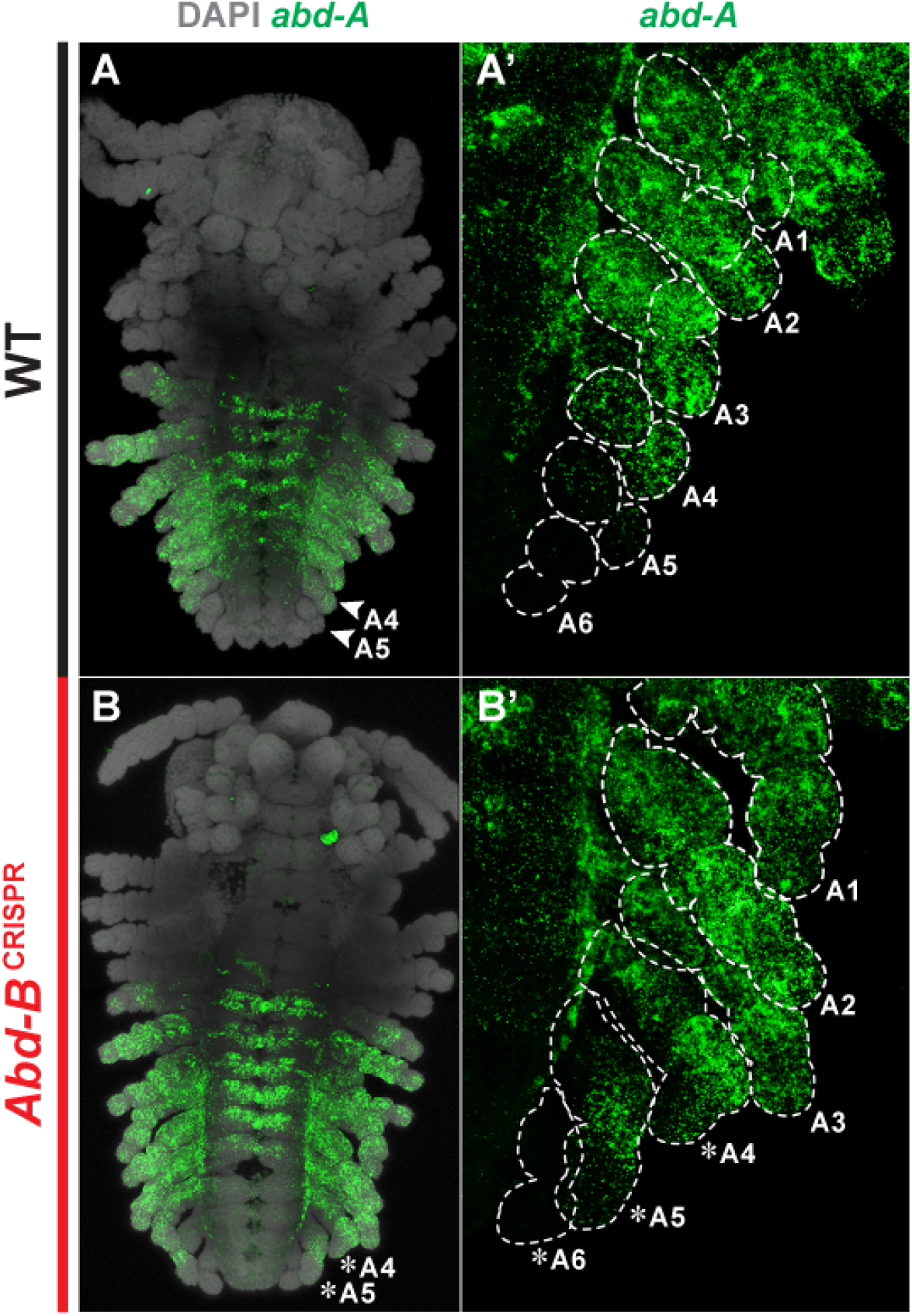
The gradated expression of *abd-A* appears unchanged by *Abd-B* knockout. A) *abd-A* expression in wildtype embryos. The expression boundary of *abd-A* delineates the distinct identities of T4/T5 forward walking legs from T6-T8 reverse jumping legs in the thorax, and A1-A3 pleopods from A4-A6 uropods in the abdomen B) *abd-A* expression in Abd-B KO embryos. A1-A6 abdominal limbs show clear transformation from bifurcated abdominal appendages to uniramous thoracic legs 6 - 7 podomeres in length. While the boundaries of *abd-A* expression are not changed with *Abd-B* KO, there appears to be some increase in the expression level. In the transformed A1 - A3 appendages, the expression of *abd-A* remained largely unchanged compared to the uninjected wildtype control embryos. The transformed A4 and A5 appendages appear to express slightly stronger levels of *abd-A* compared to their respective wildtype counterparts, though spatial restriction to the anterior domain of the limb was maintained.

#### *abd-A* expression dominantly specifies reverse jumping leg identity and combinatorially specifies swimming leg identity

*abd-A* knockout induces both posterior-to-anterior transformations and anterior-to-posterior transformations, contrary to a “posterior prevalence” effect. Loss of *abd-A* induces transformation of reverse jumping legs (T6-T8) to more anterior forward walking leg (T4/T5) identities, as well as transformation of swimming legs (A1-A3) to more posterior anchoring leg (A4-A6) identities. This result could be explained by a combinatorial logic, wherein reverse jumping leg identities are specified as a result of combined action of *Ubx* + *abd-A*, and swimming leg identities are specified by *abd-A* + *Abd-B*.

*In situ* hybridization of *abd-A* in wildtype embryos reveals that *abd-A* is expressed at high levels in the T6-T8 and A1-A3 appendages, as well as at progressively lower levels in segments A4 and A5. Additionally, within the A4 and A5 appendages, the anterior region of the limb appears to express higher levels of *abd-A* than the posterior. Despite the low levels of *abd-A* in these limbs, A4 and A5 appendages are distinctly uropod in identity. The main phenotypic difference is in their size, where the width of the A4 segment and corresponding size of uropod is significantly greater than those of A5, which are in turn greater than those of A6.

Given that *Ubx* expression expands to the posterior of the embryo as a result of *Abd-B* knockout, we wondered if the gradated *abd-A* expression was also regulated by *Abd-B*. We performed *in situ* hybridization of *abd-A* in *Abd-B* knockout embryos, and observed that the gradated expression of *abd-A* in the posterior is retained even after loss of *Abd-B*, suggesting that *Abd-B* does not repress *abd-A* expression. This result is inconsistent with a “posterior prevalence” model of *Hox* regulation, as *abd-A* expression is not derepressed as a result of *Abd-B* loss.

In *Abd-B* knockout embryos, A1-A3 appendages are transformed to reverse jumping leg identity, and appear to express both *Ubx* and *abd-A*. A possible explanation of this phenotype is that *abd-A* represses *Ubx* identity, and that segments that express both *abd-A* and *Ubx* default to the identity specified by *abd-A* alone.

However, careful analysis of the A4-A6 appendages in *Abd-B* knockouts reveals that A4 and A5 appendages appear intermediate in phenotype between the T4/5 forward walking leg and T6-T8 reverse jumping leg identities (Fig. 4B-F). The A4 appendage appeared more similar to T6-T8 reverse jumping legs than the A5 appendage, which appeared intermediate between T6-T8 and T5/T5 identity. In contrast, the A6 appendage, which lacks *abd-A* expression in both wildtype and *Abd-B* knockout embryos, appeared to exhibit near-perfect replication of the T4/T5 walking leg identity. These results reflect the stepwise decreases in *abd-A* expression along the A4-A6 appendage gradient, and are consistent with a model that *abd-A* expression works combinatorially with *Ubx* expression to set limb identity. The intermediate identity of A4 and A5 walking legs in Abd-B KO hatchlings has no clear corollary in wildtype animals, and presents an interesting example of a new combination of *Hox* gene expression generating a limb identity with morphology distinct from canonical wildtype limb morphologies.

**Fig. 4:**
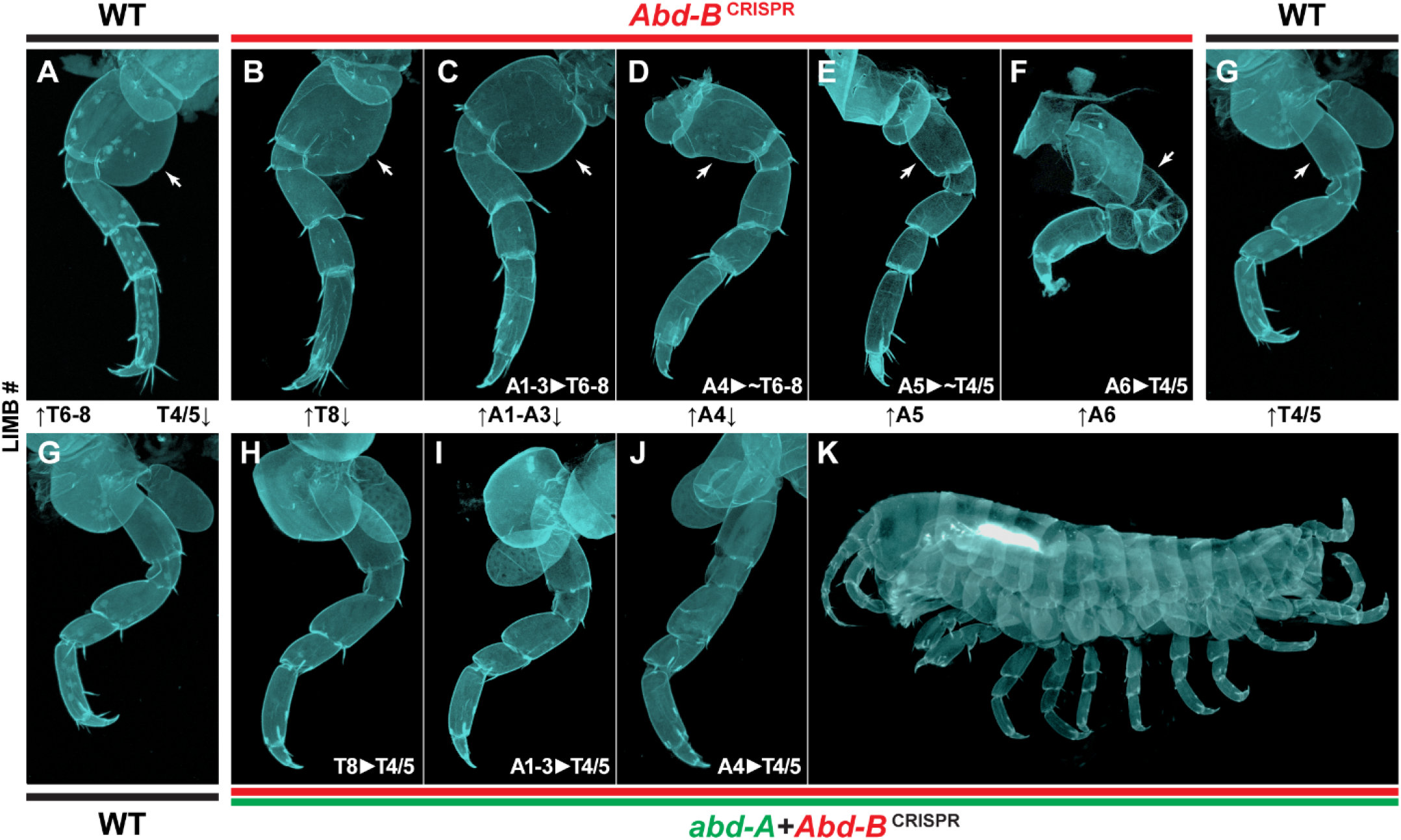
*Abd-B* knockout transformations show homeotic transformations with intermediate identities. A) Wildtype T6-8 reverse jumping leg. The enlarged basal plate is marked with an arrow. B-F) T8 and example A1-6 appendages in an *Abd-B* knockout hatchling. T6-8 appendages retain a jumping leg identity, while A1-3 appendages are transformed from a feathery swimming leg identity (see example in Supp. Fig. 2D). A4 and A5 appendages appear to show intermediate morphology between T6-8 reverse jumping and T4/5 forward walking leg identities. This is most obvious in the morphology of the basal plate (marked in each panel with an arrow), which appears enlarged in *Abd-B* knockout A4 and A5 appendages compared to *Abd-B* knockout A6 or WT T4/5. The A4 and A5 segments express lower levels of *abd-A* in a gradated fashion, while the A6 segment does not express appreciable levels of *abd-A*. Thus, the intermediate A4 and A5 appendage identities may be indicative of an additive relationship between high levels of *Ubx* and low levels of *abd-A*. G) Wildtype T4/5 forward walking leg. This panel is repeated twice for ease of comparison in each row. H-J) T8, A1-3, and A4 appendages in *abd-A* + *Abd-B* knockout hatchlings. All posterior thoracic and abdominal appendages are transformed to a T4/5 identity. This suggests that *abd-A* is necessary for the T6-8 and A1-3 jumping leg identities in *Abd-B* knockout hatchlings, and that the intermediate A4 appendage morphology is also dependent on presence of *abd-A*. K) Whole-mount *abd-A* + *Abd-B* knockout hatchling showing homeotic transformations across the posterior half of the body axis.

To further test the prediction that *Ubx* and *abd-A* must work combinatorially to produce the reverse jumping leg identity, we generated *abd-A* + *Abd-B* double-knockout embryos (Fig. 4H-K). Loss of both *abd-A* and *Abd-B* should abolish the T6-T8 reverse jumping leg identities found T6-T8, A1-A3, and intermediate A4 and A5 phenotypes in *Abd-B* knockouts. In *abd-A* + *Abd-B* double-knockout embryos, we observed that the T6-T8, A1-A3, and A4-A6 appendages all transformed towards T4/T5 forward walking leg identities. The data for this experiment is also consistent with *abd-A* having a combinatorial function along with *Ubx* to produce reverse jumping leg identity.

However, in *Ubx* + *Abd-B* knockouts, where *abd-A* expression remains (Fig. 6H-M), T6-8 and A1-3 appendages retain their reverse jumping leg identity, while the A4 and A5 identities transform towards a more intermediate identity, similar to that observed in *Abd-B* knockout alone. This suggests that, counter to previous interpretations and earlier experiments, high *abd-A* expression in the absence of *Abd-B* is sufficient to establish T6-8 reverse jumping leg identity, even without *Ubx* expression. Moreover, low levels of *abd-A* expression can generate more intermediate T4-8 morphologies on their own, without needing combinatorial input of *Ubx*. Thus, a high level of *abd-A* is both necessary and sufficient for establishing reverse jumping leg identity. Moreover, low levels of *abd-A* expression induce an intermediate limb identity between reverse jumping and forward walking legs. Finally, *abd-A* and *Abd-B* together specify swimming leg identity.

These results together indicate that the posterior limb types in *Parhyale* are specified by a combination of *Hox* cross-regulatory repression of *Ubx* by *Abd-B*, dominant action of *abd-A* over *Ubx* without the repression of *Ubx* expression, and combinatorial action of *abd-A* and *Abd-B*. Thus, crustaceans such as *Parhyale* require a combinatorial “*Hox* code” to generate their limb diversity, rather than strict posterior prevalence, revealing a substantially different mechanism from that observed for the same genes in *Drosophila*.

#### Antp and *Dfd* expression in the transformed abdominal legs of *Abd-B* KOs correspond with wildtype leg expression patterns

In addition to the *Hox* expression patterns expected to specify regional identity, *Parhyale* limbs also express more minor domains of *Hox* expression. For example, Antp protein is found in a narrow strip of non-ectodermal cells in the limbs of thoracic appendages (T4-8), while *Dfd* transcripts are expressed in the distal portion of the swimming legs (A1-3). To evaluate whether expression of these minor *Hox* domains is affected by *Hox* knockout, we examined the expression patterns of *Dfd* (using in-situ hybridization) and Antp (using immunofluorescence) in *Abd-B* KO embryos. We found that the non-ectodermal expression pattern of Antp observed in wildtype thoracic legs (T4/5 and T6-8), but not normally in abdominal appendages, is extended into the homeotically transformed abdominal legs (Fig. 5C-C’). *Dfd* is also correspondingly absent from the transformed abdominal legs that normally co-express *abd-A* and *Abd-B* (Fig. 5D-D’).

**Fig. 5:**
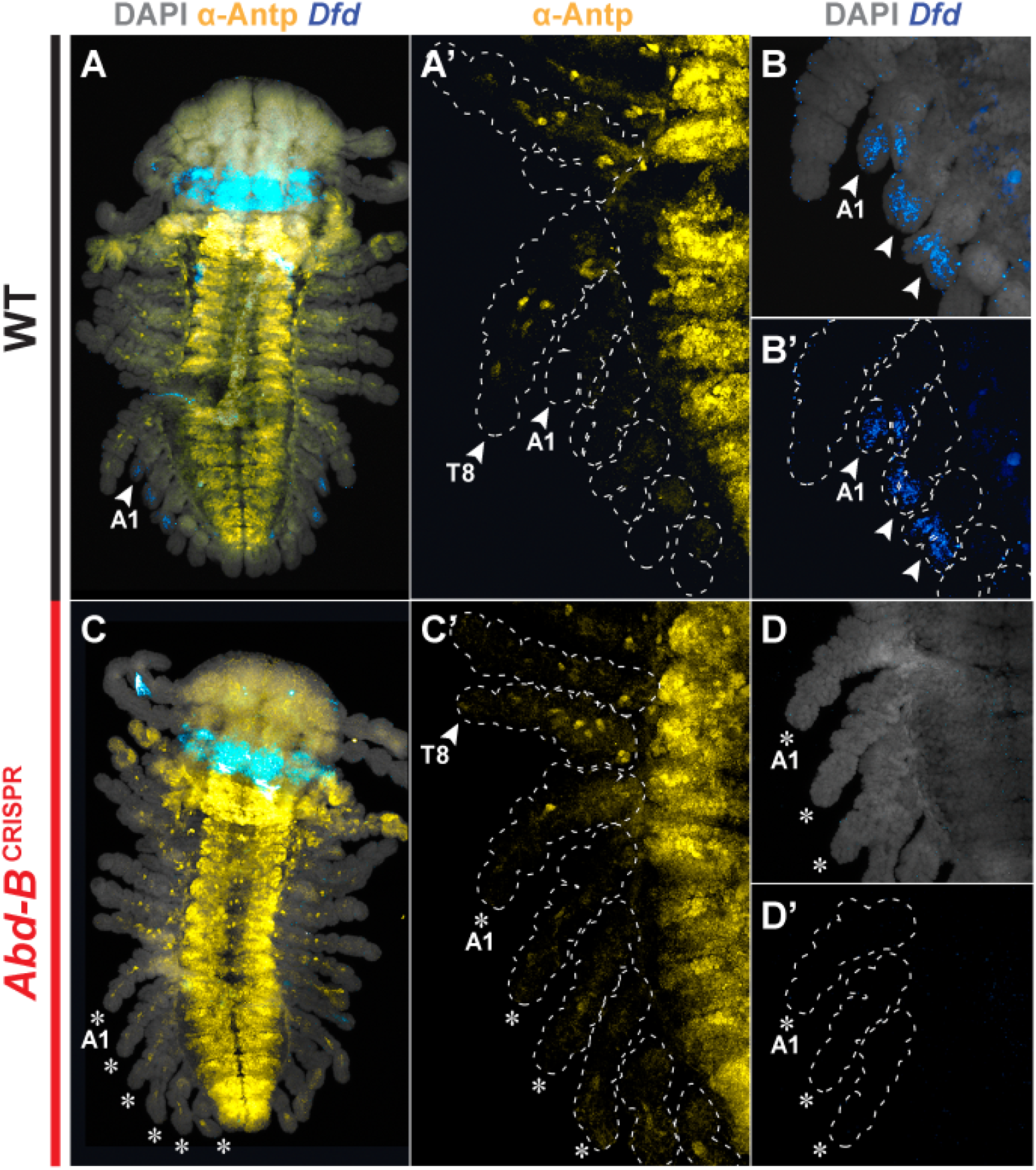
Wildtype Antp & Dfd leg patterns recapitulated by abdominal appendages transformed to legs in Abd-B KO embryos. (A-D’) Stain of stage 22 embryos by immunofluorescence (cross-reactive monoclonal ms-anti-Antp 8C11, yellow) and fluorescent in-situ hybridization (*Dfd*, blue); embryo counterstained with DAPI (gray). Arrowheads point to wildtype appendages, transformed appendages labeled with asterisks. (A-B’) Wildtype expression patterns of Antp and *Dfd*. Antp is expressed in the T1 maxilliped and T-3 claws and in a limited, non-ectodermal pattern in the remaining T3-8 thoracic legs. Antp is not expressed in the abdominal appendages (B). While *Dfd* is expressed primarily in the maxillary appendages Mn and Mx1 (A), this study focuses on its ancillary expression in nascent wildtype pleopods (B-B’). (C-D’) In *Abd-B* KO embryos, the same non-ectodermal Antp patterning exhibited by T4-8 thoracic legs is ectopically expressed in the A1-6 abdominal appendages that are clearly transformed to a thoracic-like leg identity (C-C’). Furthermore, the A1-3 expression of *Dfd* in wildtype pleopods (but not thoracic legs) is absent from the transformed legs (D-D’). As is shown in Figure 2, *Ubx* is expressed throughout the *Abd-B* KO transformed abdomen of this specific embryo (not shown).

These data describe the effects of *Abd-B* knockout on the expression of all *Hox* genes normally found in the wildtype thorax and abdomen of Parhyale. Our data demonstrate that the *Hox* expression underlying each of the limbs transformed by *Abd-B* KO corresponds to the unique combination of *Hox* genes expressed by the respective wildtype limb each is transformed toward. These data suggest that minor *Hox* expression domains are activated, directly or indirectly, by the expression of particular combinations of *Hox* genes: the non-ectodermal Antp domain appears to be activated either by *Ubx* expression of *Ubx* + *abd-A* expression, while the swimming leg-specific *Dfd* expression domain requires the expression of both *abd-A* and *Abd-B*. These results reveal that some of the downstream targets of combinatorial *Hox* expression in *Parhyale* may include other *Hox* genes.

### Cross-regulatory interactions among *Hox* genes *Scr, Ubx, abd-A*, and *Abd-B* explain non-canonical appendage transformations

The above observation of interactions among *Abd-B, abd-A*, and *Ubx* in the specification of limb identities strongly suggests that combinatorial interactions on target genes control aspects of appendage morphology in *Parhyale*. In order to further explore such interactions, we systematically examined the transformed limb morphologies for the remaining *Ubx, abd-A*, and *Abd-B* double KO combinations as well as the triple KO of all three. Each set of transformations was accompanied by a post-KO expression analysis to uncover the shifts in expression domains of other *Hox* genes that may be regulated by cross-regulatory interactions and to examine the underlying combinations of *Hox* genes putatively responsible for each transformed identity.

#### Simultaneous KO of Abd-B and Ubx transforms the posterior-most appendages toward a maxilliped fate

Upon knockout of *Abd-B, Ubx* expression expands to the posterior of the embryo, demonstrating that *Abd-B* represses *Ubx*. In these mutants, the correlation between the expansion of *Ubx* expression and the non-contiguous transformation of abdominal appendages to thoracic appendages suggests that *Ubx* is responsible for specifying the newly transformed identities. To validate that *Ubx* expression is directly responsible for these transformations, we performed simultaneous knockout of *Abd-B* and *Ubx* (Fig. 6). Strikingly, the double KO of *Ubx* + *Abd-B* results in the posterior-most abdominal appendages taking on maxillary features of the anterior-most thoracic identity: the shorter, branching gnathal T1 maxilliped (Mxp) (Figure 5A-C; G-I). In some cases, the A6 appendage underwent complete transformation to a maxilliped identity (Fig. 5C). A1-3 appendages, however, continued to exhibit the same reverse jumping leg transformation as with *Abd-B* KO alone. This demonstrates that *Ubx* is necessary not only for the development of forward walking leg identity in normal development, but also for the homeotically transformed limbs in *Abd-B* knockouts.

**Fig. 6:**
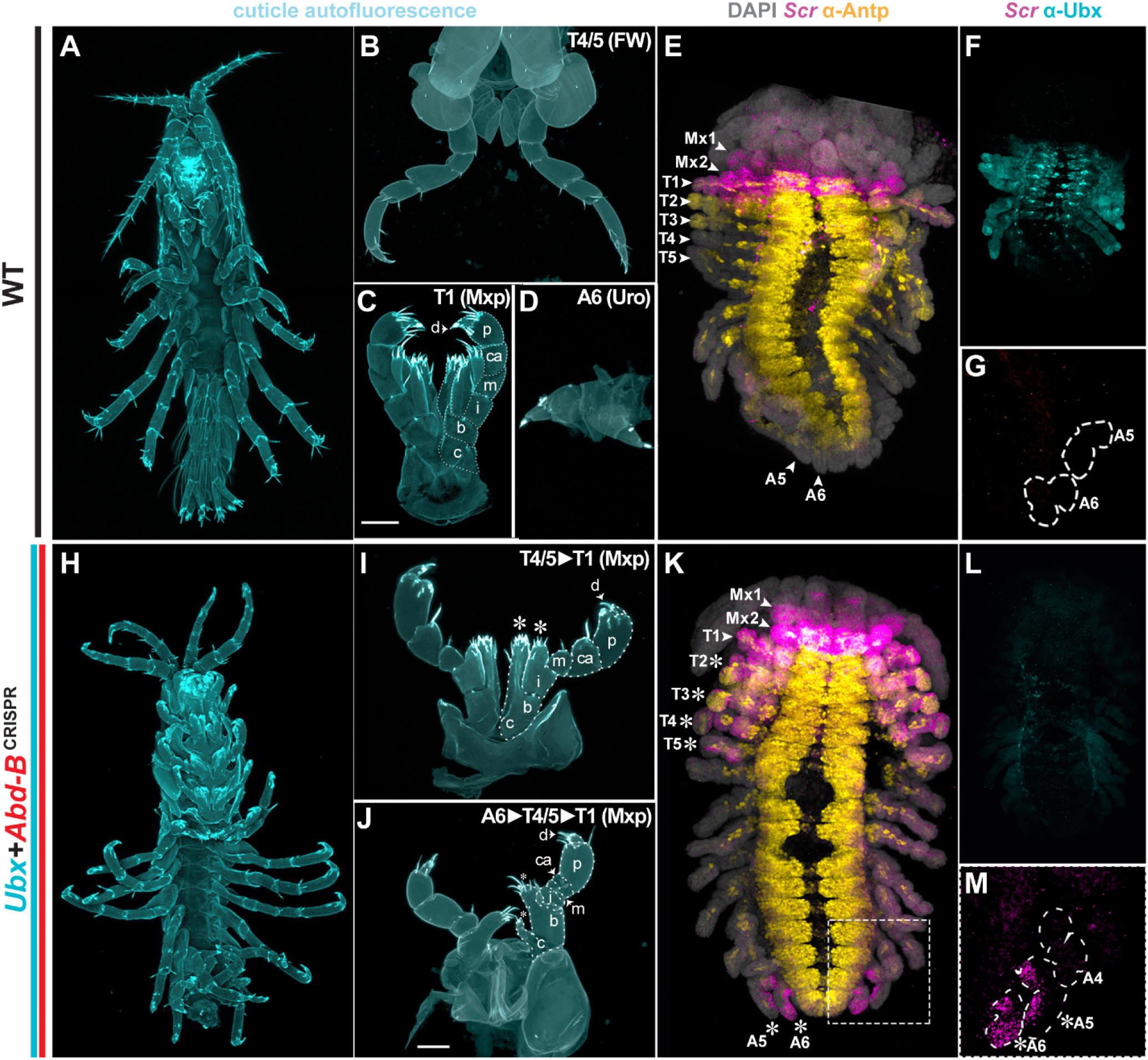
Simultaneous knockout of the posterior Hox genes *Ubx* and *Abd-B* results in the ectopic expression of Scr and a corresponding maxilliped identity. A-D) Wildtype hatchling and limbs; H-J) *Ubx* + *Abd-B* hatchling and limbs; Wildtype (E-G) and *Ubx* + *Abd-B* (K-M) S22 emb ryo stained using anti-Ph-Ubx (cyan), anti-Antp (8C11, yellow), and *in situ* hybridization of *Scr* (pink). H-J) *Ubx* + *Abd-B* double knockout transforms the posterior-most walking leg (the A6 appendage, which lacks *abd-A*) towards a maxilliped fate (note that single KO of *Ubx* transformants show no transformation in the A6 uropod identity). The *abd-A* expressing appendages T6-8 (reverse jumping legs) and A1-3 (also transformed to reverse jumping legs by single *Abd-B* knockout) exhibit the jumping leg phenotype exhibited in Abd-B KO alone. The intermediate walking-jumping legs exhibited by A4 and A5 in the single KO of Abd-B (which have lowered levels of anteriorly-restricted abd-A) continue to exhibit their respective Abd-B KO identities as well. As is exhibited by the single KO of Ubx, the anterior thoracic appendages T2/T3 (claws) and T4/T5 (forward walking legs) are also transformed towards the maxilliped identity. When paired with the simultaneous use of an additional Ubx guide (Table ?), our results yield T4/T5 maxilliped transformations that are much more complete than has previously been demonstrated. Right panels show the corresponding embryonic expression of Scr, Antp, and UBX in wildtype (D-F) vs. Ubx + Abd-B KO (J-L) animals. Multi-label fluorescent staining panels show the wildtype and post-knockout expression of Scr (in-situ hybridization) and ANTP and UBX (immunofluorescence) show that the transformation towards a maxilliped identity is paired with loss of Ubx (relative its ectopic expansion upon Abd-B knockout) and gain of Scr. 40x zoom shows that the Scr expression in A6 is expressed fully throughout the entire developing limb, whereas its expression in A5---which expresses low-levels of abd-A in its anterior compartment---appears to be limited to the posterior compartment.

Previous studies functionally demonstrated the role of *Ubx* in selecting leg vs. maxillary identities at the anterior boundary of *Ubx* expression (Liubicich et al. 2009; Pavlopoulos et al. 2009; Martin et al. 2016). This study is the first, to our knowledge, to demonstrate the potential for maxillary transformation beyond the posterior boundary of normal *Ubx* expression. Consistent with the single KO of *Ubx*, double *Ubx* + *Abd-B* KO transforms the claws (T2/3) and forward walking legs (T4/5), but not reverse jumping legs (T6-8), towards the shorter, branching gnathal phenotype of the T1 maxilliped (Figure 1D). While previous experiments achieved only partial transformation towards a maxilliped identity in T4/5 walking limbs, the simultaneous use of multiple *Ubx* guides allowed us to achieve complete maxilliped transformation of all T2-T5 appendages.

#### Double knockout of *Ubx* and *Abd-B* induces ectopic expression of *Scr*

In Parhyale, the maxilliped identity is associated with the expression of *Scr* and *Antp*. Previous work (Martin et al., 2015) functionally demonstrated that the loss of *Scr* leads to the corresponding loss of the maxillary branches and an overall lengthening of the primary endopod. We performed a post-knockout expression analysis of *Scr* and Antp on *Ubx* + *Abd-B* double knockout embryos and clearly observed ectopic expression of *Scr*, but not Antp, in the transformed limb primordia of the anterior thorax and posterior abdomen (Fig. 6). This result demonstrates the cross-regulatory repression of *Scr* by the more posterior *Hox* genes *Ubx* and *Abd-B*. Moreover, this result indicates that *Scr* specifies maxilliped identity. The absence of a posterior shift in the Antp expression domain suggests that regulation of the posterior boundary of Antp is not dependent on the three posterior *Hox* genes (*Ubx, abd-A*, and *Abd-B*), and may occur independently from *Hox* cross-regulatory mechanisms, as is true for *abd-A*.

#### *Ubx* + *abd-A* + *Abd-B* triple knockout transforms all appendages of the thorax and abdomen to the maxilliped fate

The ectopic *Scr* expression and the associated transformation to maxilliped identity in *Ubx* + *Abd-B* double knockout is only observed in regions of the thorax and abdomen that lack the expression of *abd-A*. This suggests that the expression of *Scr* and the associated maxilliped identity may also be repressed by *abd-A*. To test this cross-regulatory interaction and examine *abd-A* function in these processes, we carried out the simultaneous triple knockout of *Ubx* + *Abd-B* + *abd-A*. Strikingly, but not surprisingly, triple-knockout hatchlings exhibit the maxilliped identity for all thoracic and abdominal appendages (Fig. 7).

**Fig. 7:**
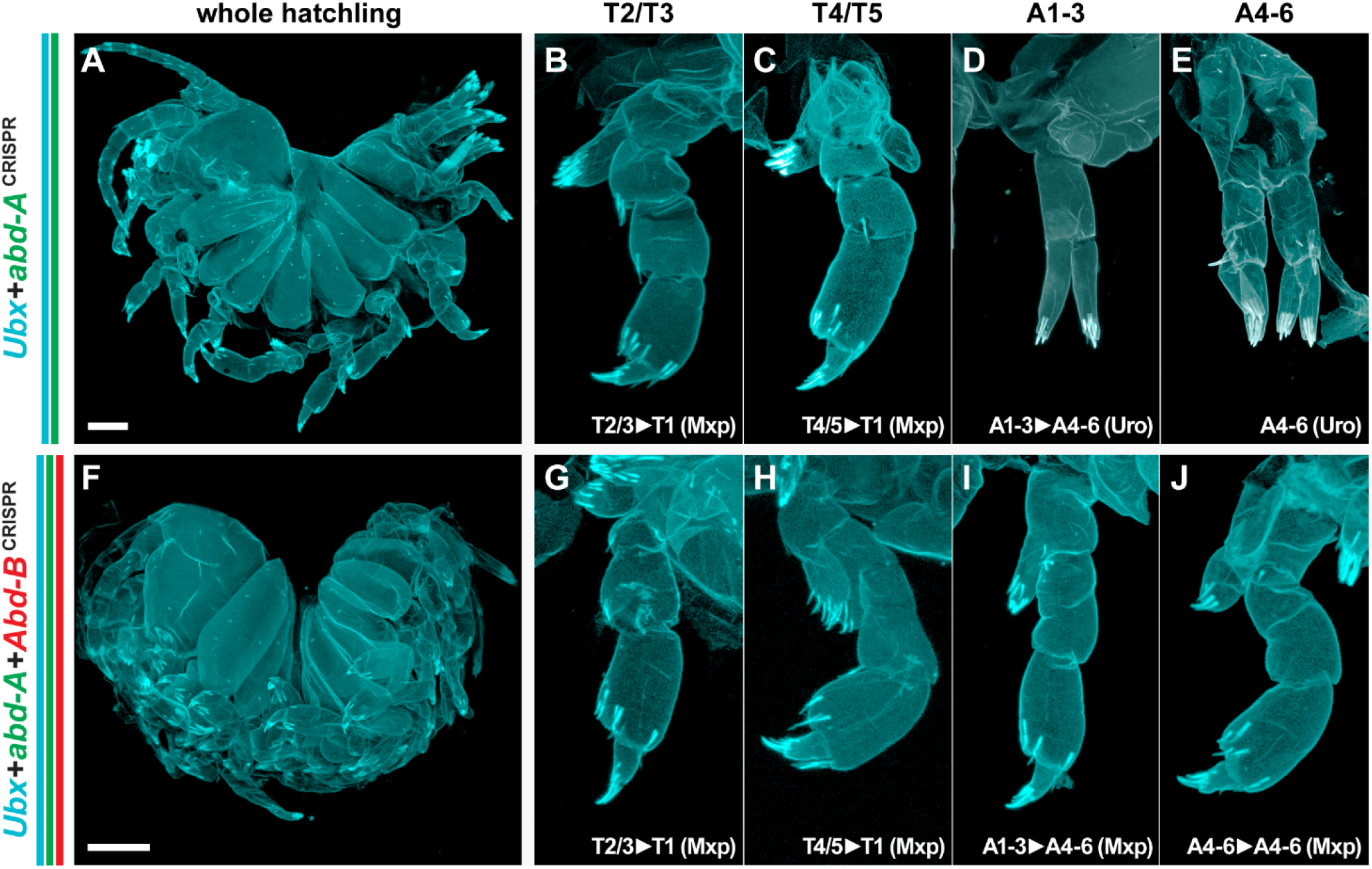
Triple knockout of the posterior *Hox* genes induces broad homeosis to maxilliped identity. A) *Ubx* + *abd-A* CRISPR mutant hatchling exhibiting strong homeoses across the body axis. B-E) *Ubx* + *abd-A* CRISPR mutant limbs. Thoracic appendages, such as T2/3 and T4/5, are transformed to T1/Mxp identity, as predicted by repression of *Scr* by *Ubx* and *abd-A*. Abdominal appendages A1-3 are transformed to A4-6 uropod identity in the absence of *abd-A*. F) *Ubx* + *abd-A* + *Abd-B* CRISPR mutant hatchling exhibiting homeotic transformations. G-J) *Ubx* + *abd-A* + *Abd-B* CRISPR mutant limbs. In comparison to *Ubx* + *abd-A* mutants, these mutants have transformations of A1-3 and A4-6 appendages to T1/Mxp identity. Thus, all thoracic and abdominal appendages are transformed to a T1/Mxp-like identity in the absence of posterior *Hox* genes.

**Fig. 8:**
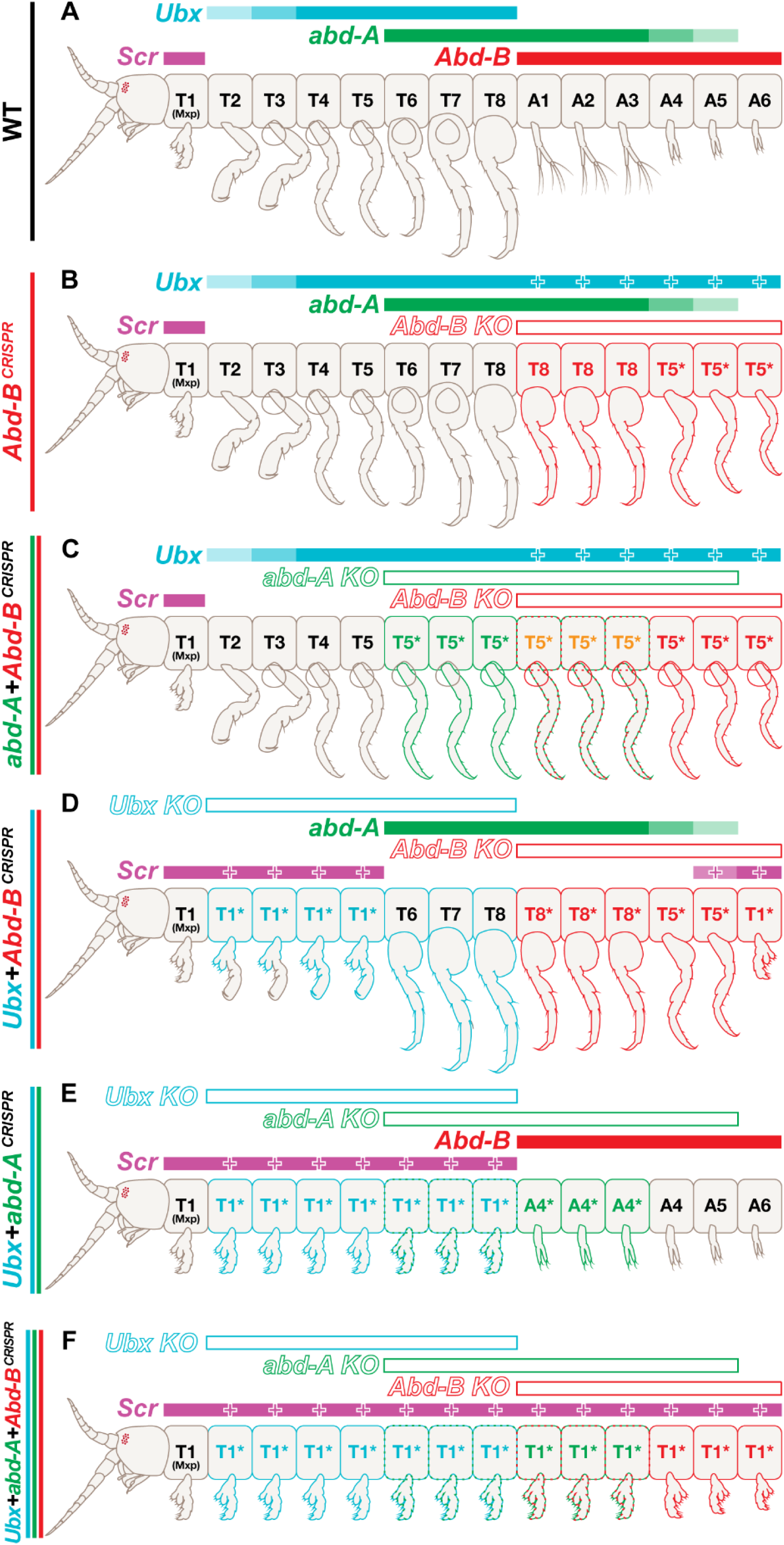
Summary of double- and triple-*Hox* knockout phenotypes. A) Wildtype *Parhyale* thorax and abdomen and wildtype *Hox* expression. B) *Abd-B* knockout exhibiting transformations of A1-3 to T6-8 identity and A4-6 to T4/5 identity. This non-contiguous transformation is explained by expansion of *Ubx* expression into the posterior. C) *abd-A* + *Abd-B* double knockout exhibiting transformations of T6-8, A1-3, and A4-6 to T4/5 forward walking leg identity. *abd-A* is necessary for the jumping leg identity, *Abd-B* is necessary for the anchoring leg identity, and *abd-A* + *Abd-B* together specify the swimming leg identity. D) *Ubx* + *Abd-B* double knockout exhibiting transformations of T2-3, T4-5, and A4-6 to T1/Mxp identity. Loss of both *Ubx* and *Abd-B* results in expansion of *Scr* expression, revealing that *Ubx* and *Abd-B* both repress *Scr. Scr* expression is not expanded in segments expressing high levels of *abd-A*. E) *Ubx* + *abd-A* double knockout exhibiting transformations of T2-3, T4-5, and T6-8 to T1/Mxp identity, as well as transformation of A1-3 to A4-6 anchoring leg identity. F) *Ubx* + *abd-A* + *Abd-B* triple knockout exhibiting transformations of all thoracic and abdominal appendages to T1/Mxp identity.

**Fig. 9:**
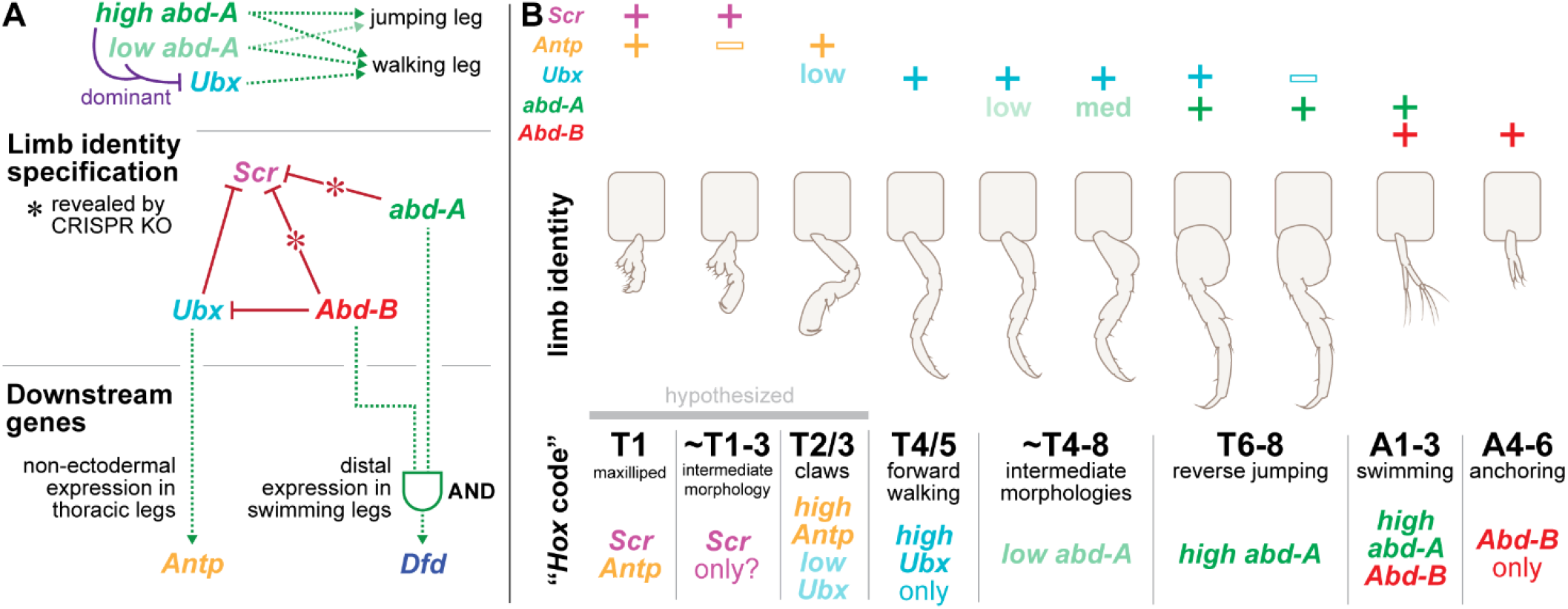
Summary of *Hox* cross-regulatory interactions and the proposed *Parhyale Hox* code. A) *Hox* cross-regulatory interactions revealed by CRISPR experiments. High levels of *abd-A* appear to act dominantly to *Ubx*, whereas low levels of *abd-A* appear to work additively with *Ubx*, as evidenced by intermediate phenotypes in *Abd-B* knockout A4 and A5 limbs. *Scr* is repressed by *Ubx, abd-A*, and *Abd-B*, as revealed by individual and combinatorial knockouts of each of these three genes. *abd-A* and *Abd-B* suppression are not normally observable in single knockouts, but become revealed upon multiple *Hox* knockouts. In addition, *Abd-B* represses *Ubx* to set the posterior boundary of *Ubx* expression. Downstream of these *Hox* genes, *Antp* is activated by *Ubx* expression both in wildtype limbs and in ectopic T4-8 identity limbs found in *Abd-B* knockout animals. *abd-A* and *Abd-B* are both required in order for *Dfd* expression and swimming leg identity to develop, suggesting a true combinatorial input is upstream of *Dfd* expression in that region. B) The “*Hox*” code of the *Parhyale* thorax and abdomen. Different combinations of *Hox* genes result in different limb morphologies. Some morphologies, such as the *Scr*-only T1-3 intermediate appendage or the T4-8 intermediate morphologies in *Abd-B* knockout abdominal appendages, are not found in wildtype animals. *abd-A* and *Abd-B* work combinatorially to specify swimming leg identity. We hypothesize that combinatorial interactions between *Scr* and *Antp*, as well as *Antp* and *Ubx*, are responsible for specifying the T1/Mxp and T2/3 claw identities.

In order to examine the complete set of *Ubx, abd-A*, and *Abd-B* KO combinations, we performed double knockout of *Ubx* and *abd-A*. As our *“Hox* code” model would predict, all thoracic limbs were transformed to maxillipeds, and swimming legs (A1-A3) were transformed into anchoring leg identities (A4-6)(Fig. 7). Thus, by performing all combination of knockouts of the three posterior *Hox* genes, we have been able to establish a posterior “*Hox* code” for the *Parhyale* body plan, and to uncover cross-regulatory interactions between *Hox* genes in this organism.

## Discussion

Studies across diverse arthropod taxa have revealed that *Hox* genes are critical to specifying regional identity. In some clades, such as crustaceans, differences in *Hox* expression domains between species appear to strongly correlate with differences in the organism’s body plan. For example, loss of *Ubx* and gain of *Scr* in thoracic appendages appears to result in the evolution of maxilliped identity in the crustacean trunk (Abzhanov and Kaufman, 1999; Averof and Patel, 1997; Fritsch and Richter, 2017; Liubicich et al., 2009; Pavlopoulos et al., 2009).

These studies have been informed, in part, by the large body of *Hox* regulatory studies in *Drosophila*, from which useful models such as the “posterior prevalence” model have been derived (Struhl and White, 1985). However, as our work has demonstrated, the elegance and simplicity of posterior prevalence is not sufficient to explain how crustacean *Hox* genes function in patterning the body. Indeed, even within *Drosophila*, the rules of posterior prevalence do not seem to apply to the anterior *Hox* genes, and work in other organisms suggests that protostomes do not have a universal directionality to their *Hox* cross-regulatory interactions (Durston, 2012; Gehring et al., 2009).

Here we have examined the cross-regulatory interactions and combinatorial effects of the Hox genes Scr, Ubx, abd-A, and Abd-B in establishing appendage identities in the Parhyale thorax and abdomen. We find that a simple posterior prevalence model is also not sufficient to explain the transformations observed upon *Hox* knockout, and that a “*Hox* code” model best explains the role of *Hox* genes in the formation of *Parhyale*’s diverse range of appendage types.

We have examined the appendage transformations for all single, double, and triple KO combinations of the posterior *Parhyale Hox* genes *Ubx, abd-A*, and *Abd-B* and determined the patterns of *Hox* expression associated with each mutant background. We uncovered a number of cross-regulatory interactions that overall explain the pattern of appendage identity in mutant and wild type animals. The *Hox* expression patterns underlying each specific transformed appendage type correspond with morphologically similar wildtype appendages and can be explained by altered expression domains of *Hox* genes affected by the respective *Hox* gene knockouts. Furthermore, appendage transformations that exhibited novel intermediate identities have patterns of *Hox* expression distinct from any wildtype pattern. Interestingly, the novel combinations of features in these intermediate transformed appendages reflect aspects of the morphologies associated with wild type *Hox* codes.

### Some *Parhyale Hox* genes establish posterior boundaries through cross-regulation

In this study, we demonstrate that *Parhyale Hox* genes utilize cross-regulation to establish wildtype expression boundaries. For example, we show that *Abd-B* represses *Ubx* and is essential for setting the posterior boundary of *Ubx* expression. Knockout of *Abd-B* permits a posterior expansion of *Ubx* throughout the abdomen and that this shift in expression induces corresponding homeotic transformations, explaining the previous non-contiguous transformations observed in previous work.

We also demonstrate that *Ubx, abd-A*, and *Abd-B* all suppress *Scr* expression, and are essential for establishing its posterior boundary. Upon combinatorial knockout of *Ubx, abd-A*, and *Abd-B*, limbs that would be expected to have no *Hox* expression (in the absence of *Hox* cross-regulation) begin to express *Scr*. Some of these interactions could only be observed through knockouts of multiple *Hox* genes; for example, double knockout of *Ubx* and *Abd-B* results in a hatchling where all thoracic and abdominal appendages – aside from those retaining *abd-A* expression – are transformed to a T1/Mxp-like identity. These transformed T1/Mxp appendages also express *Scr*, but do not appear to express high levels of *Antp*. Triple knockout (*Ubx* + *abd-A* + *Abd-B*) reveals that *abd-A* also represses *Scr*; all thoracic and abdominal appendages in those hatchlings are transformed into T1/Mxp-like identity. Thus, each of the three posterior *Hox* genes is capable of repressing *Scr* expression.

### *Hox* cross-regulation can also activate *Hox* expression

In addition to observing instances of derepression and homeotic transformation upon loss of *Hox* expression, we also observed changes to ancillary *Hox* expression patterns upon transformation. For example, Antp is normally expressed in non-ectodermal cells of some thoracic limbs (T4-T8), observed as low levels of diffuse expression, as well as individual high-Antp expressing cells in the middle of those thoracic limbs. Upon homeotic transformation of abdominal limbs to thoracic T4-T8 limbs in *Abd-B* knockouts, we also observed expansion of Antp expression into the non-ectodermal cells of the transformed limbs. As another example, *Dfd* is normally expressed in the distal ends of swimming legs (A1-3, pleopods) specified by combined *abd-A* and *Abd-B* expression. Upon loss of *abd-A* and transformation of A1-3 appendages to anchoring leg (A4-6, uropod) identity, *Dfd* expression was lost.

While it is not possible with our data to determine whether these interactions are a result of direct regulation of one *Hox* gene upon another, these results do reveal that some of the downstream targets of *Hox* identity specification may include other *Hox* genes. Upon homeotic transformation, markers other than the identity-specifying genes also exhibit transformed expression. Thus, *Hox* cross-regulation in *Parhyale* appears to potentially involve both repressive and activating mechanisms.

### “Posterior prevalence” is not sufficient to explain *Parhyale Hox* expression patterns and limb identities

The presence of cross-regulatory repression in some *Parhyale Hox* genes resembles a “posterior prevalence” model of *Hox* regulation. For both *Ubx* and *Scr*, posterior *Hox* genes are essential to repress more anterior genes. However, our data reveal that posterior prevalence alone is not sufficient to explain a number of limb identities and *Hox* expression patterns in mutants. In particular, by examining which transformations and shifts in expression were *not* observed upon different combinations of *Hox* knockouts, we reveal that strict posterior prevalence cannot fully explain *Hox* cross-regulatory interactions in *Parhyale*.

First, none of the combinations of *Hox* knockouts created homeotic transformations towards swimming leg identities (A1-3, pleopods). Limbs exhibiting the pleopod identity were only observed in segments where *abd-A* and *Abd-B* were co-expressed. Previous work illustrated that CRISPR knockout of *Dfd*, another gene expressed in swimming legs, did not induce homeotic transformations of these appendages. Loss of function of *Ubx* also did not result in additional pleopod emergence. Thus, the establishment of pleopod identity requires multiple *Hox* genes, rather than the dominant “posterior prevalence” effect of a single *Hox* gene.

Second, the overall pattern of *abd-A* expression was not perturbed upon CRISPR knockout of either *Ubx, Abd-B*, or both *Ubx* and *Abd-B*. We did not observe anterior or posterior shifts in *abd-A* expression in either knockout, contrary to the effects of *Abd-B* knockout on *Ubx* expression. This suggests that the anterior and posterior boundaries of *abd-A* expression are not regulated by *Hox* cross-regulatory mechanisms emerging from its two overlapping neighbors, *Ubx* and *Abd-B*.

Finally, the posterior boundary of *Antp* also did not appear to be perturbed as a result of loss of *Ubx* expression, its posterior neighbor. Loss of *Abd-B* in the posterior also did not result in derepression of *Antp*, while it did result in derepression of *Scr*. This suggests that the *Antp* posterior boundary is also not regulated by “posterior prevalence”-like *Hox* cross-regulatory repression. Given that CRISPR mutagenesis of *Scr* results in a transformation of T1 appendages towards T2/3 morphology, it is possible that *Scr* partially represses *Antp*. Such an interaction would explain the lack of expanded *Antp* expression in the posterior of the embryo upon loss of *Ubx* and *Abd-B*. This proposed mechanism would also explain why T2/3 appendages appear to become reduced in response to loss of *Ubx*: *Scr* in *Ubx* knockouts becomes de-repressed, partially expanding into the T2/3 appendages to create transformations towards a mixed T1-T2/3 identity. If this hypothesis is true, it would represent an example of an anterior *Hox* gene repressing a more posterior *Hox* gene in a more similar flavor to the “anterior prevalence” of some *Drosophila Hox* genes.

Thus, while some *Parhyale Hox* genes require repressive cross-regulation in order to establish proper wildtype expression boundaries, a strict posterior prevalence model is not sufficient to explain the wildtype pattern of *Parhyale Hox* genes. Moreover, the presence of limb identities that require the simultaneous expression of multiple *Hox* genes suggests that combinatorial logic must also play a role in patterning the *Parhyale* body plan.

### *abd-A* interacts dominantly with *Ubx* and combinatorially with *Abd-B* to specify limb identity

Our data suggest that, rather than a posterior prevalence model, a “*Hox* code” model may better explain the specification of limb identities in the *Parhyale* body plan. In particular, the *Hox* gene *abd-A* appears to display dominant effects with *Ubx* and combinatorial effects with *Abd-B*. Joint expression of *Ubx* and *abd-A* results in a dominant effect and a jumping leg identity, whereas combinatorial expression of *abd-A* and *Abd-B* is required for swimming leg identity.

In our knockout experiments, we observed that swimming legs (A1-3, pleopods) were only formed when *abd-A* and *Abd-B* expression co-occurred within the same limb segment. This result demonstrates that swimming leg identity is necessarily combinatorial. Moreover, we also observed that loss of *abd-A* also resulted in loss of *Dfd* expression in the A1-3 appendages, indicating that this specific expression pattern is dependent on both *abd-A* and *Abd-B* expression. Thus, a combination of two *Hox* genes is required not only for the morphology of the swimming leg identities, but at least one downstream *Hox* target gene appears to require combinatorial input by multiple *Hox* genes.

Previous work, including individual knockouts and expression data, suggested that the reverse jumping leg (T6-8, reverse pereopod) identity required combinatorial input by *abd-A* and *Ubx*. Given that jumping legs express both *abd-A* and *Ubx*, and loss of *abd-A* in single CRISPR knockout resulted in loss of jumping leg identity, it was previously reasonable to argue that jumping leg identity might require both genes. However, combinatorial knockout of *Ubx* and *Abd-B* resulted in the development of limbs exclusively expressing *abd-A*, namely T6-8 and A1-3 in knockout embryos. These limbs, contrary to a combinatorial model, appeared to develop clear jumping leg identity.

While T6-8 and A1-3 appendages, which express high levels of *abd-A*, retained jumping leg identities, the appendages A4 and A5 did not develop clear jumping leg identity. These two limbs express lower levels of *abd-A* in a stepwise gradient towards the posterior of the embryo. In both *Abd-B* single knockout and *Ubx* + *Abd-B* double knockout embryos, A4 and A5 developed an intermediate morphology between jumping and walking legs. This suggests that *abd-A* alone can specify reverse jumping leg identity and that the previous interpretation of *Ubx* and *abd-A* working combinatorially to specify jumping leg identity is incorrect.

Moreover, the intermediate phenotypes produced in *Abd-B* and *Ubx* + *Abd-B* knockouts suggest that *abd-A* interacts in a dose-dependent fashion to specify leg identity. Low levels of *abd-A* alone or in combination with *Ubx* produce morphology similar to forward walking legs with the increased basal plate width. This suggests that both *Ubx* and *abd-A* activate a uniramous walking leg program, but that high levels of *abd-A* can drive a forward walking leg towards the jumping leg identity. Low levels of *abd-A* activate both the forward walking leg identity, as well as weaker activation of the jumping leg identity, resulting in an intermediate phenotype.

The suggestion that *abd-A* function may be dose-dependent is consistent with previous work, wherein varying levels of heat shock misexpression of *Ubx* resulted in variable limb morphologies, including limbs of intermediate phenotypes not identical to normal wildtype limbs. Moreover, within the *Parhyale* body plan, several *Hox* genes exhibit variable expression levels. *Ubx*, for example, is weakly expressed in the claws (T2-3, chelipeds), and this low level of expression appears to be essential for normal claw morphology. Thus, *Hox* genes in crustaceans may utilize not only Boolean categories of *Hox* expression, but also variable levels of *Hox* expression, to achieve morphological diversity.

Recent work has begun to reveal that combinatorial *Hox* expression, in particular combinatorial *abd-A* expression, appears to play a role in numerous other groups of arthropods. Expression data from several other malacostracan crustacean species including *Procambarus fallax* (crayfish), *Mysidium columbiae* (mysid), and *Porcellio scaber* (woodlouse) show that the distinction and relative numbering of limb subtype in both the thorax and abdomen correspond to their respective overlap with *abd-A* (Abzhanov and Kaufman, 2000; Martin et al., 2015). This suggests that shifts in the boundaries of *abd-A* expression may have provided a modular mechanism for macroevolutionary change in crustaceans.

The combinatorial ability of *abd-A* can also be found in hexapods. Although *Ubx* and *abd-A* suppress the formation of limbs in most insects, a number of basal hexapods retain their abdominal appendages. In the collembolan *Orchesella cincta* (springtail), combined expression of *Oc-Ubx* and *Oc-abd-A* specifies formation of a specialized stabilizing appendage that is different from the springtube specified by *Oc-Ubx* or the leaping organ specified by *Oc-abd-A* alone (Konopova and Akam, 2014). siRNA knockdown of *Oc-Ubx* results in the transformation of the A3 stabilizing appendage toward the more posterior leaping organ of A4 that expresses *Oc-abd-A* alone. Rather than following the expected prediction of posterior prevalence, this outcome parallels the *abd-A* KO transformation the *Parhyale* abdomen, supporting the potential for combinatorial function of *abd-A* and other *Hox* genes in diverse arthropod taxa.

### *Scr* expression changes may drive maxilliped evolution

Our results indicate that posterior *Hox* genes repress *Scr*, and that loss of posterior *Hox* expression results in the expansion of T1/maxilliped morphology further into the thorax. This phenotype is reminiscent of the macroevolutionary gain (and loss) of maxillipeds with the retraction (or expansion) of the anterior expression boundary of Ubx (Averof and Patel, 1997). Previous studies have demonstrated through loss-of-function (Liubicich et al., 2009; Martin et al., 2015) and gain-of-function (Pavlopoulos et al., 2009) experiments that *Ubx* expression establishes the posterior boundary of maxilliped identity. Our study is the first to demonstrate that maxilliped identity can develop in more posterior thoracic and abdominal regions in the absence of other *Hox* expression.

Previous work has also demonstrated that *Scr* function is essential for maxilliped identity. CRISPR mutagenesis of *Scr* results in loss of maxilliped identity, and heat shock overexpression-induced loss of maxillipeds appears to stem from repression of wildtype *Scr* by ectopic *Ubx* expression. Using our combined knockout experiments, such as *Ubx* + *Abd-B*, we are able to demonstrate that *Scr* positively regulates maxilliped identity; loss of *Ubx* and *Abd-B* results in expanded *Scr* expression and homeotic transformations to maxillipeds. These results suggest that, in addition to a posterior shift in *Ubx* expression, crustaceans that expand or retract maxilliped identities should likely also exhibit reciprocal changes to *Scr* expression. Thus, the evolution of maxilliped identity among crustaceans could be an example of how *Hox* cross-regulatory mechanisms are able to drive macroevolutionary changes.

## Conclusion

*Hox* genes have sparked considerable attention since their discovery, primarily due to the striking homeotic transformations observed when their expression or function are perturbed. Embryos are patterned not by a series of *Hox* genes in isolation, however. Axial regionalization is contingent to some degree upon the cross-regulatory interactions among *Hox* genes. These can occur on many levels, including pre-translational refinement of one another’s expression boundaries and post-translational selection of identity in regions where multiple Hox domains overlap (Noro et al., 2011; Slattery et al., 2011). In addition, *Hox* genes may synergistically or competitively co-regulate downstream pathways, where the genetic interactions among *Hox* genes modulate phenotype by fine-tuning the establishment of one another’s boundaries through cross-regulatory interactions.

By studying post-transformational *Hox* expression in *Parhyale Hox* knockouts, we have revealed numerous cross-regulatory interactions between the *Parhyale Hox* genes. Our data suggest that cross-regulatory and combinatorial interactions are both crucial to limb identity specification in *Parhyale*. We have shown that a classical posterior prevalence model is not sufficient to predict and understand the appendage transformations in *Parhyale*. Rather, we propose a more modular “*Hox* code” regulates limb identities in *Parhyale*, wherein co-expression of *Hox* genes is required for certain limb identities. This study demonstrates that interactions among *Hox* genes provide a mechanism for the diversification of appendage morphologies and the modularization of integrated appendage arrangements over evolutionary time.

## Materials and Methods

### CRISPR/Cas9 Somatic Mutagenesis

Recombinant Cas9-NLS protein (QB3, UC Berkeley) was combined with purified single-guide RNA (sgRNA) to form a ribonucleoprotein complex (RNP) and delivered directly by microinjection. Details of RNP assembly and Parhyale-specific genome editing techniques developed by our lab are visually described in (Farboud et al.). CRISPR guides Abd-B#1, abd-A#1, and Ubx#1 are described in (Martin et al., 2016). Guides new to this study were designed using Geneious software (Biomatters, version R9 (Kearse et al., 2012)) and generated synthetically by Synthego (Redwood City, CA). We found ordering synthetic RNA to be a significant time-saver and cost-effective, and, in our hands, the editing efficiency and survival rates of Synthego sgRNA were equivalent to or superior to that generated by in vitro transcription. See Table 1 for guide sequences, rates of survival and efficiency, and reference to published *Parhyale Hox* gene sequences.

Approximately 40-60 picoliter of the final RNP injection mix (4 - 8 uM sgRNA (150-200 ng/uL) + 2 uM Cas9 (333ng/uL) + 0.05% phenol red for visualization) was microinjected into one-cell (or both blastomeres of two-cell) embryos following the published protocol (Rehm et al., 2009e). Injected embryos, along with batch/age-matched uninjected controls, were cultured at 26°C (12h day-night cycles) to hatching (for phenotypic and genetic analysis) or sacrificed midway through embryogenesis for downstream embryonic Hox expression analyses.

### Staining and Imaging

The expression patterns for multiple Hox genes were visualized in parallel by in-situ hybridization (*Dfd, Scr, abd-A*, and *Abd-B*) and/or immunofluorescence (Ubx and Antp) following modified versions of the published protocols (Rehm et al., 2009c; Rehm et al., 2009d). In-situ probes are described in (Serano et al., 2016) and labeled with either Digoxigenin (DIG) or Dinitrophenol (DNP) conjugated UTPs. When performed in parallel, *Hox*-specific and anti-hapten primaries were added concurrently after hybridization. In-situ hybridization reactions were visualized either fluorescently with AlexaFluor secondaries in parallel with immunofluorescence or enzymatically using FastRed (Sigma F4648). Antibodies are as follows. Hox-specific primaries: polyclonal Rat-anti-Ph-Ubx (1:5000) (Liubicich et al., 2009), cross-reactive anti-Antp at 1:40 (DSHB monoclonal, 8C11, source?), and cross-reactive anti-Ubx/Abd-A) at 1:20 (DSHB monoclonal FP6.87, Kelsh et al.(1994)). Anti-hapten primaries: sheep αDIG (Roche) and/or Rabbit anti-DNP (Thermo Fisher) or αDIG-AP. Secondary antibodies were used at 1:500 and included donkey-anti-rabbit AlexaFluor-488 and/or donkey-anti-rat AlexaFluor-647 (Jackson ImmunoResearch) and/or donkey anti-Sheep AlexaFluor-555 (ThermoFisher Scientific). In-situ antibody scheme modified from (Ronshaugen and Levine, 2004).

Expression panels were performed on G0 CRISPR/Cas9 KO Parhyale embryos along with age-matched uninjected control from the same brood batches. Embryos intended for downstream expression analyses were cultured for approximately 115 - 125h (S21/S22) for in-situ hybridization and 120 - 132h (Stage 21) for immunofluorescence. We found these age ranges optimal for balancing embryonic development that has advanced enough for the fate of the nascent limbs to become distinguishable but before the deposition of cuticle in later stages becomes problematic. Embryonic staging is based on (Browne et al., 2005). Embryos were dissected out of their membranes in 3.2% paraformaldehyde (PFA) in filter sterilized seawater or in a 9:1:1 ratio of PEM buffer, 10xPBS, and 32% PFA following published methods (Rehm et al., 2009b) and fixed for a total of 20 minutes (immunofluorescence) or 40 minutes (in-situ hybridization). A combination of immunofluorescence (UBX and ANTP) and/or in-situ hybridization (dfd, Scr, abd-A, and Abd-B) was performed against multiple Hox genes in parallel

Parhyale hatchlings were sacrificed at 0 - 3 days post hatching and fixed in 3.2% paraformaldehyde as described above for approximately 30 minutes. Fixed hatchlings were placed in glycerol in a glass bottom dish to image the overall arrangement of appendages. Individual appendages were then carefully removed using sharpened tungsten needles.

Parhyale are direct developers with the appendage morphology of hatchlings reflecting the adult morphology. Hatchlings appendages and stained embryos were mounted with 70% glycerol on glass slides, and imaged using a LSM 780 scanning laser confocal (Zeiss) using 10x (entire hatchlings) 20x (individual hatchling legs and entire embryos) or 40x magnification (embryo appendages). LSM files were processed using Volocity software (Perkin-Elmer) and individual channels overlaid in Photoshop (Adobe). Schematic representations made in Illustrator (Adobe).

## Acknowledgements

We would like to thank Matt Ronshaugen for his invaluable edits and improvements to the manuscript, Panda Kreidler for animal care and tool making, and Heather Bruce for being willing to lend a helpful hand. EJ would like to thank Kevin Chung for finishing my experiments when she was unable. We would also like to thank the staff at Synthego (Redwood City, CA) for technical assistance and hand-delivered sgRNA guides.

## Competing Interests

The authors declare no competing interests.

## Author Contributions

NHP and EJ conceived of the project. EJ carried out staining experiments, microscopy, and analysis. EJ, KC, and SR performed CRISPR/Cas9 mutagenesis. EJ wrote the manuscript. DAS revised the manuscript and provided additional images and interpretations of the data. NHP edited the manuscript and provided feedback to EJ and DAS.

## Notes

### Competing Interest Statement

The authors have declared no competing interest.

