## Supplementary figures and images for "Combinatorial interactions of *Hox* genes establish appendage diversity of the amphipod crustacean *Parhyale hawaiensis*"

### Supplemental Figures

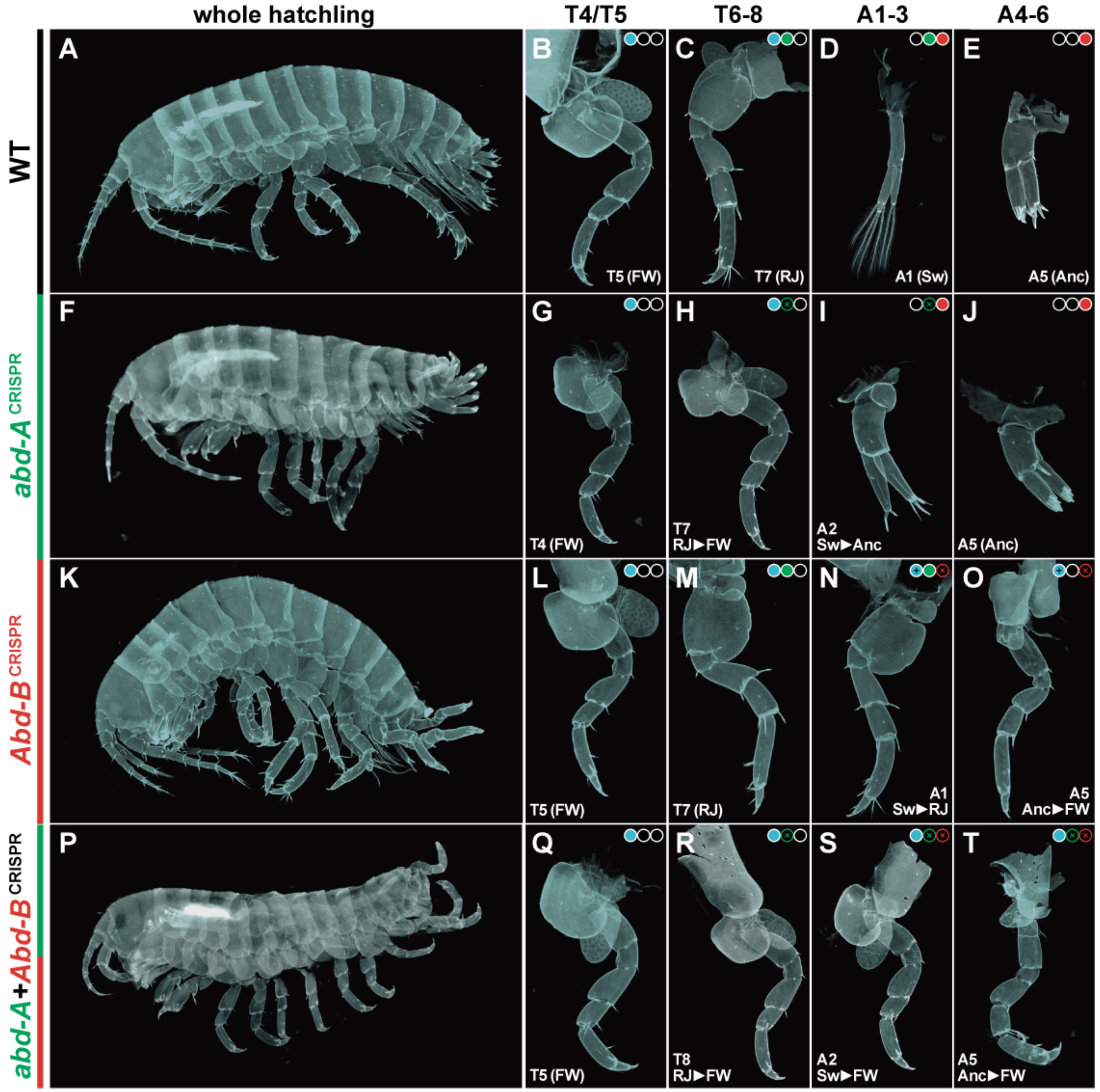


**Supp. Fig. 1: Summaries of *abd-A*, *Abd-B*, and *abd-A* + *Abd-B* mutant phenotypes.**
